# The inhibitory cascade model and evolution in segmentally organized tissues

**DOI:** 10.1101/2025.10.20.683466

**Authors:** Benjamin M Auerbach, Charles C Roseman

## Abstract

The inhibitory cascade model (ICM) of morphogenesis is an effort to link development to the production of variation, which can influence evolutionary trajectories. The ICM proposes that serially developing features, such as molar teeth, is governed by the relative magnitudes of one activating and one inhibiting developmental process. The statistical expectations of the ICM are typically expressed and analyzed on a first-element standardized scale and seem to be a good predictor of molar proportions. However, the ICM has been applied to traits that *occur* in series but do not *develop* in sequence and still recovers as good a fit as when applied to serially developing traits. Such an undiscriminating result raises questions about whether the fit of the ICM is an artifact of standardization. The mathematical rendition of the ICM do not correspond with the verbal descriptions of the developmental argument. Applying our novel re-articluation of the ICM to biological, non-biological, and simulated data, we demonstrate that the apparent goodness of fit of the ICM to many biological systems is an artifact of scaling correlated values with a common denominator. There is little evidence supporting the ICM at the developmental, variational, or evolutionary levels.

## 1 Introduction

A central aim of evolutionary developmental biology is to build connections between the development of organisms, the manifestation of variation in populations of organisms, and the evolution of phenotypes (Hall 2012). This development-variation-evolution chain involves addressing two challenges. The first involves building models of how variation in developmental process leads to distributions of phenotypic variation in populations (e.g., Arthur 2004; Stansfield and Parsons 2024). The second challenge entails modeling how this generation of variation via development enables or inhibits evolutionary transitions across different time scales (e.g., Wagner 1988; Hansen 1997; Hansen and Houle 2008; Jablonski 2020). At short time scales, the evolution of phenotype depends on variation among individuals arising from differential developmental outcomes. Over longer time scales, patterns of macroevolution can be shaped by limitations imposed by developmental and physical constraints and internal stabilizing selection (Schmalhausen 1949; Hallgrímsson et al. 2012, Hallgrímsson et al. 2023) on the ranges of phenotypic variation.

Surmounting the explanatory challenges presented by these multi-level and multiscale issues is particularly difficult because phenotypic variation provides little information about the identity and operation of the developmental processes that produce such variation (Hallgrímsson et al. 2009). The variance of any continuous complex trait is the product of developmental processes that conduct many genetic and environmental influences of small effect into phenotypic variation (Boyle et al. 2017; Liu et al. 2019). Among the efforts to link development to phenotypic variance within an evolutionary framework are the association of modular phenotypic patterns to genetic effects (e.g., pleiotropy) (Cheverud 2007), as well as documenting genomic mechanisms, such as shared regulation, that have evidence for accelerated evolution (Davidson and Erwin 2006; Senevirathne et al. 2025).

Almost two decades ago, Kavanagh et al. (2007) proposed the inhibitory cascade model (ICM) as one solution to address these challenges in modeling the developmentvariation-evolution theoretical chain in the case of mammalian molar teeth. Kavanagh et al. described the mechanistic basis of the ICM as a ratcheting mechanism governed by the relative strengths of inhibiting and activating factors during the development of molar teeth in series from front to back in the tooth row. A relative overabundance of activating factors would result in a ratcheting increase in molar size from anterior to posterior along the tooth row. In contrast, a relative overabundance of inhibiting factors would result in a ratcheting down of the molar size along the tooth row. In a case in which the activating and inhibiting processes were of the same magnitude, the size of the teeth would be even across the tooth row. This ratcheting mechanism would produce a distinct pattern of the sizes of the second and third molars standardized by the size of the first molar (Figure 1).

**Figure 1.**
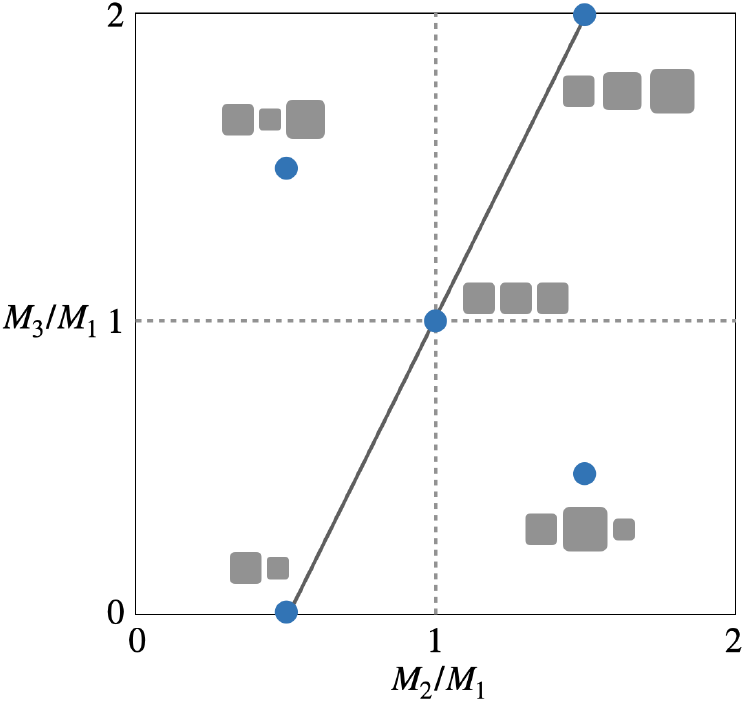
Morphospace showing mean expectations (the line) of ratio proportionality expressed in the ICM as shown in Kavanagh et al. (2007), where M2/M1 is the ratio of the second element and the first element, and M3/M1 the ratio of the third and first element. Molar size patterns in the upper left and lower right quadrants fall outside the expectations in the ICM.

The initial empirical investigations of the ICM reported what appeared to be strong support for the model. Kavanagh et al. (2007) demonstrated in murine models that the eventual proportional sizes of the teeth could be altered via manipulation of the early lamina in ways that comported well with the hypothesized activating and inhibiting mechanisms. They further reported that the linear relationship in relative tooth sizes was a good fit to the expectations of the ICM (the line with intercept -1 and slope 2 in Figure 1). That is, the relative sizes of the second and third molars standardized by the size of the first molar appeared to be roughly in line with ICM at both the within-population and macroevolutionary scales (Kavanagh et al. 2007; Polly 2007; Labonne et al. 2012; Grieco et al. 2013; Halliday and Goswami 2013).

These early successes of the ICM inspired a fluorescence of studies, mostly at the macroevolutionary scale (e.g., Asahara 2013; Young et al. 2015; Hardin 2020; Bermúdez de Castro et al. 2021; Nikolic et al. 2023). Although the many empirical applications of the ICM appear to support the predictions of the ICM linear model, several studies of molar teeth have yielded results that deviate from ICM expectations (Polly 2007; Asahara 2013; Bernal et al. 2013; Hlusko et al. 2016; Roseman and Delezene 2019; Vitek et al. 2020; Boughner et al. 2021; Bermúdez de Castro et al. 2021). Roseman and Delezene (2019) compared predicted primate molar size variation and covariation based on the ICM and direct estimates of variance and covariance, finding the ICM failed to predict the variational properties of molar teeth in seven out of eight species. They cautioned against relying on the ICM as a sole explanatory model and––salient to our study here––that departures from ICM expectations need more context to be useful in hypothesis testing of evolutionary and developmental outcomes. The ICM, furthermore, has a mixed precision in predicting within-species character covariance and poor performance in interspecific comparisons (Bernal et al. 2013; Halliday and Goswami 2013; Vitek et al. 2020). Reasons cited for this lack of fit range from a lack of understanding how evolutionary divergences from the ICM might have occurred in crown species to incomplete knowledge about the genotype-phenotype map.

Applications to the relative sizes of limb elements, vertebrae, and body segments also demonstrated fits to the ICM expectations even though these features of organisms do not develop in series in the same way as molar teeth (e.g, limb segmentation development; see Bénazet and Zeller 2009; Towers et al. 2012). That a model developed to explain the relative sizes of traits that occur and develop in series appears to be a good fit to traits that *occur* in series but do not *develop* in series is a sign that there may be a problem with the developmental argument of the model. Moreover, the expectations of the ICM concerning the proportional sizes of the elements are often accepted as good characterization of empirical results at multiple levels of organization, but the variational expectations on the raw measurement scale are seldom met. This suggests that the apparent good fit of the model on the proportional scale to that on the raw scale might be an artifact of standardization.

Ratios of correlated characteristics have statistical properties that mask sources of variation and thus are not amenable to clear evolutionary analysis, as information about whether the numerator, denominator, or both traits in the ratio are changing when evolution occurs is hidden in the calculation of the ratio (Atchley et al. 1976; Atchley and Anderson 1978). Moreover, standardization using a common denominator can make even random noise arising from measurement error appear as non-trivial correlations among traits even if all traits in the system are statistically independent. Because of these statistical properties of ratios, positive correlations among serialized traits arising from genuine biological processes that do not resemble those proposed by the ICM can yield variational and macroevolutionary patterns bearing a strong qualitative resemblance to those expected by the ICM. Characteristics that occur in serial may exhibit a decaying pattern of covariance between adjacent and nonadjacent elements simply because of proximity. Such patterns may alone explain the fit of the ICM to a variety of serial structures even if a ratchet-like activation-inhibition developmental process is absent.

In this paper, we reexamine the theoretical and empirical bases of the ICM. We demonstrate that much of the apparent goodness-of-fit of the ICM is attributable to an artifact arising from the choice of standardization. To reach our conclusion, we examine the developmental basis of tooth size variation, translate the model into a causal analysis framework, and derive expectations for the ICM for both within-population covariation within and among traits and between-species macroevolutionary divergence on the raw and proportional scales. We also demonstrate how differences in variational properties of serially occurring traits can lead to expectations that fit and depart from the ICM. We emphasize the importance of measurement theory and understanding the effects of standardization and scale in the construction of the development-variation-evolution theoretical chain. Through our analyses, we highlight the importance of testing fundamental statistical assumptions when building models and open new avenues for future research into the development and evolution of serialized traits.

### 1.1 Assessing the ICM and its applications

The ICM is applied across three interrelated scales: developmental, variational, and evolutionary. On the developmental scale, the model is applied to predict the results of interventions into developmental processes, both natural and experimental, within the lifespan of organisms (e.g., murine models in Kavanagh et al. 2007). The ICM also yields predictions about variation in cross-sectional groups of organisms at similar developmental stages in a population. Lastly, on evolutionary time scales, the ICM provides comparative predictions about which evolutionary pathways are more likely at various time scales. While multiple studies have marshaled results that are broadly supportive of the ICM at the developmental, variational, and evolutionary scales, we identify here several pressing shortcomings of the model.

#### 1.1.1 Re-evaluating the developmental argument for the ICM

Kavanagh et al. (2007) operationalized a general view of developmental regulation by activation-inhibition that has roots in morphogenic wave models first proposed by Turing (1952). Morphogenic wave models propose that tissue growth and differentiation during morphogenesis are driven by spatially and temporally structured gradients in the concentrations of different morphogenic signaling factors. Under the ICM, then, the initiation and growth of serially arranged tissues would be controlled by a balance of activating and inhibiting factors expressed in an order that precisely timed the formation, promotion of differentiation and growth, and inhibition of growth in tissues based on signals from adjacent tissues that develop earlier in the growth series. The variation in size and shape would therefore be the result of minor perturbations in only two factors responsible for the timing and promotion of tissue development.

Kavanagh et al. (2007) noted that little was known at the time about the specific developmental mechanisms that might be responsible for the activation-inhibition predicted by the ICM for molars (*cf*. Balic and Thesleff 2015; Polly 2015; Balic 2019; Hermans et al. 2021). Kavanagh et al. argued, however, that their experimental manipulation of the serial development of molar teeth in dental laminae implicated a mechanism in which the relative abundance of inhibiting and activating factors along gradients occurred similar to Turing’s model. They grew murine molars *in vivo* and *in vitro*, and in the latter left the dental lamina intact or separated the ectodermal placodes of the second molar in the lamina from the developing first molar. When the lamina was intact, the development of the second molar was delayed, and separation of the dental lamina resulted in accelerated development of the second molar and a larger ultimate size. Kavanagh et al. concluded that this was evidence for the effects of decreased inhibition when the lamina was separated.

Since their publication, multiple studies have added to the field’s understanding of the specific molecular and histological bases of tooth formation and development. Kavanagh et al. 2007 focused on the expression of *Bmp4*, which was shown be suppressed in adjacent dental lamina in explants, most likely by the secreted protein ectodin. Introduction of BMP4 via an implanted bead rescued delayed formation of a dental placode and bud, thus giving Kavanagh et al. reason to think that tooth formation that occurs earlier could inhibit the activation of adjacent teeth formation that occurs later.

More recent studies indicate, however, that *Bmp4* is downstream of the Wnt/*β*-catenin pathway in the mesenchyme of the developing alveolus, which is also associated with epithelial cell migrations (probably of multipotent stem cells) that give rise to the invagination of ectodermal placodes before their transition to the bud stage (see Hermans et al. 2021). The formation of the initial lamina is subject to the coexpression of multiple signaling molecules and genes, including *Msx, Dlx, Lhx* family genes in the presence of PAX and RUNX2, as well as transcription factors like SP6 (Rhodes et al. 2021). These and other transcription factors, such as SOX2, are expressed in the extension of the epithelium to form successive teeth, but again this is an activation signal and not an inhibition signal (Juuri et al. 2013; Yu et al. 2015). In fact, the role of transcription factors and signaling molecules (expressed in adjacent dental placodes) in the dental identity and size of adjacent dentition is not known. Factors associated with both shape and size of teeth are likely located in the surrounding mesenchyme in the dental crypt and not the adjacent dental lamina (Bei 2009; Balic and Thesleff 2015). To this point, most signaling for the initiation of tooth formation is controlled by the transient primary enamel knot signaling center within the developing tooth (Jernvall et al. 1994), but this only *activates* dental formation and does not control the overall size of teeth or the length of development.

Thus, the *in vitro* experiments reported by Kavanagh et al. (2007) as support for an inhibition cascade in molar development may have excluded important sources of signaling and regulation from adjacent tissues not included in the explants. Kavanagh et al. at best show that the expression of factors in *M*_1_ delay the initiation of *M*_2_ growth, but do not inhibit its growth once it starts. Changing the developmental timing of *M*_2_ and *M*_3_ does affect their ultimate size in the Kavanagh et al. experiment, but this is an *in vitro* experiment lacking the interactions with the adjacent mesenchyme. As Kavanagh et al. note, leaving the surrounding mesenchyme in place would have stunted tooth development in culture. That missing mesenchyme is *essential* in regulating dental development, mediated through signaling molecule and miRNA interactions with multipotent stem cells that give rise to the enamel knots (Yu et al. 2015). Furthermore, the idea that the activating influence determines dental size ignores the many downstream tissue interactions between developing dentin and enamel that yield both tooth shape and size. Most of the regulation of dentin and enamel differentiation in teeth comes from interactions between the dental epithelium and the mesenchyme, and not from signals from adjacent teeth (Balic and Thesleff 2015). Once the cap stage is reached, the Wnt/*β*-catenin pathway continues to control the development of cusp size, and *Shh* and *Sostdc1* are involved in regulating the intercusp distances (and thus tooth shape). Their proteins are secreted by and regulated in secondary enamel knots that derive from the primary enamel knot (see Hermans et al. 2021). Thus, dental size is not the product of simple regulation of activation timing for formation, but instead is a function of constant interactions with adjacent mesenchyme and signaling centers within developing teeth.

#### 1.1.2 Within-population variation

This lack of support for a clear ICM-like mechanism at the developmental level is compounded by the rejection of the ICM expectations for within-population variance and covariance within and among elements occurring in series. Humans (Bermúdez de Castro et al. 2021), an assortment of fossil mammals (Vitek et al. 2020), and primates (Roseman and Delezene 2019) all displayed patterns of variance, covariance, and molar size that were discordant to ICM predictions when applied to within-species samples. In the case of Vitek et al. and Roseman and Delezene, these departures from ICM expectations are particularly convincing because the predictions were precise quantitative expressions of the model as opposed to generic applications of regression or qualitative interpretations of variation.

The within-population variational expectations of the ICM includes the variance of *M*_3_, the covariance of *M*_3_ with the other elements, and that the covariance matrix among all elements should be singular, or nearly so, in that one dimension in the morphospace defined by tooth size. With one exception (*M. mulatta*, Roseman and Delezene 2019), no species conforms to the variational expectations involving *M*_3_. Moreover, no species exhibits a singular covariance matrix or a lack of conditional covariation between molars.

#### 1.1.3 Application of the ICM in comparative contexts

Polly (2007) expanded the comparative analysis of Kavanagh et al. (2007) predictions for tooth proportions to a comparison of molars across mammals. He argued that the ICM is, for the most part, a good predictor of among-species distributions of proportional molar size. Polly’s analysis formed the template for other papers that displayed varying degrees of fit to the ICM (Grieco et al. 2013; Hlusko et al. 2016; Hardin 2020). For instance, Carter and Worthington (2016) applied the model to tooth size in species of primates. They concluded that relationships among the proportional sizes of teeth were in qualitative agreement with the ICM across different samples, further arguing that highly conserved ICM-compatible developmental mechanisms structured tooth evolution across the primate order (some 60my to 80my).

Yet, not all studies of the macroevolutionary distribution of molar proportions support ICM expectations. Studies by Asahara (2013) in canids, Evans et al. (2016) in fossil hominins, Bermúdez de Castro et al. (2021) in modern humans, and Schroer and Wood (2015), Carter and Worthington (2016), Hlusko et al. (2016), and Machado et al. (2023) in primates, all reported departures. For example, Schroer and Wood found few exceptions to the expectations of the ICM in Catarrhine monkeys, though they noted that the pattern of molar sizes in *Paranthropous* hominins did not fit well. Notably, Carter and Worthington showed that the ICM predicts that there should be no covariance between the first and third elements when the second element is included as a mediator and decisively rejected this observation. Likewise, Hlusko et al. (2016) found the fit of the ICM to be so dissatisfying that they sought out entirely new means of expressing dental variation in comparative contexts. These departures from ICM expectations are typically not taken as cause for rejecting the ICM *in toto*, as researchers appeal to unknown developmental processes (e.g., Bermúdez de Castro et al. 2021, p. 1178) and natural selection acting on non-IC generated variation to drive dietary adaptation (e.g., Asahara 2013).

### 1.2 Lack of fit & the specter of artifact

In our review of studies incorporating the ICM, we find a contrast: macroevolutionary studies find broad support for it but the within-population variation scale does not. With the addition of the concerns we presented about its developmental basis, this contrast raises the question of how a particular developmental process constrains or enables evolutionary trajectories if it is not the predominant cause of the disposition of within-population variation. Two possibilities come to mind: 1) developmental constraint may be acting on higher moments of the distribution, thus imparting an evolutionary bias without it being evident in the variation within populations; or 2) the way in which the macroevolutionary expectations are evaluated makes them appear to fit the model better. There is an important clue, though, that an artifact might be responsible for the pattern. The ICM developmental model explicitly applies to adjacent segmental traits that form in a serial developmental sequence. But its expectations fit segmental structures that do not develop this way. For example, Young et al. (2015) reported that ICM evolutionary expectations were well supported in limb and somite segmentation in vertebrates, though neither forms like molars (see Towers et al. 2012; Maroto et al. 2012). Likewise, Nikolic et al. (2023) applied the ICM to segmentation in invertebrates and vertebrates to argue that activation-inhibition is a common mechanism by which serial trait size patterns are determined during development. That is, they used phenotypic variation to infer a developmental process where it is not previously described.

This strategy of inferring a process using pattern alone comes with risks, especially when applied to the proportional scale typical of ICM-structured studies. For one, developmental processes are underdetermined by variation such that many different developmental systems may channel genetic and environmental influences into the same pattern of variation (i.e., equifinality) (Cheverud 2007; Gompel and Prud’homme 2009; Hallgrímsson et al. 2009).

Standardizing the second and third element by the first element in the system, however, is a more pressing concern in the context of the ICM. No previous study has expressed the linear model expectations of the ICM on the proportional scale or taken the statistical properties of ratios of correlated variables (e.g., Atchley et al. 1976; Atchley 1978) into account. A particular concern here is the way in which ratios of positively correlated variables with a common positively correlated denominator can lead to strong spurious correlations (Atchley and Anderson 1978). Moreover, choice of measurement and data transformations can motivate results in counter-intuitive ways and produce artifactual results that can difficult to detect. Thus, we focus on the mathematical expectations of the ICM for the remainder of this paper, as it is apparent that the inconsistencies in prior results we presented above may be the collective result of a mathematical artifact.

## 2 Materials and Methods

Here we describe: a) a recasting of the ICM in terms of a path model; b) a description of data used in this analysis; and c) a description of the statistical and simulation methods used to analyze the data.

### 2.1 The mathematical inhibitory cascade model

Kavanagh et al. (2007) establish the ICM expectations for the sizes of the *k*^*th*^ elements of serialized traits in an individual *i* as determined by each *k* element divided by the size of the first element 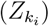. 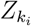 can be expressed as a function of the order of appearance in the series (*k*_*i*_) and the effect on size of what Kavanagh et al. (2007) call activating factors (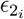 or *a* in Kavanagh et al.), and the effect on size of inhibiting factors (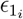 or *i* in Kavanagh et al.). This can be expressed as:

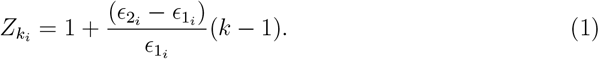

We adopt the notation of Morrissey (2015) in the place of Kavanagh et al. (2007) to avoid using the term *i* as a value other than an index of individuals and to make clear a distinction between exogenous developmental inputs into a system (*ϵ*_1_ and *ϵ*_2_), the endogenous morphological features of the system (the element sizes: *M*_1_, *M*_2_, and *M*_3_), and the relationship of exogenous developmental and endogenous morphological influences on fitness (*W* ). We use the rules of causal modeling (Wright 1921, 1934; Simon 1957; Selltiz 1959; Blalock. 1964; Morrissey 2015) to convert these systems of equations into a path model relating the exogenous developmental inputs to the endogenous morphological outcomes and then on to fitness. Figure 2 expresses the ICM in path diagram form on the raw measurement scale as opposed to the first element standardized scale given in Eq. 1.

**Figure 2.**
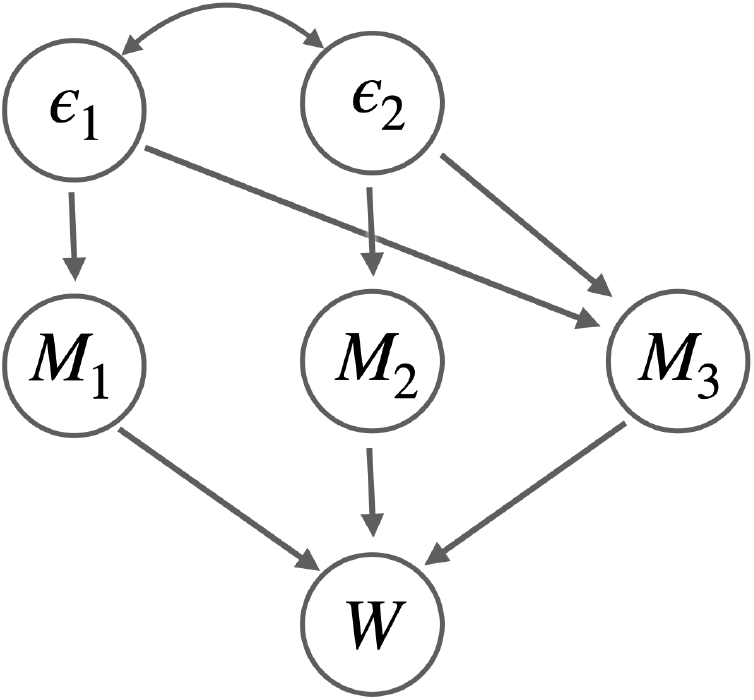
Path diagram of the ICM in the idiom of Morrissey (2015) relating the two exogenous developmental variables (*ϵ*_1_ and *ϵ*_2_, corresponding to *i* and *a*, respectively, in Kavanagh et al. 2007) to the size of the endogenous morphological features in a serially growing system (*M*_1_, *M*_2_, and *M*_3_), and then fitness (*W* ).

The equivalent of Eq. 1 on the raw measurement scale is given by

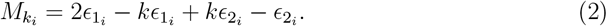

Expressing each of the first three elements in a system in terms of Eq. 2 gives us 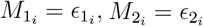, and 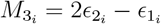.

### 2.2 Data and analyses

We test whether values of serially occurring traits (or events) expressed as proportions of the first value in the series can approximate the predictions of the ICM as a result of any process that results in correlated outcomes. To assess this, we use race times from the 2024 Formula 1 season (https://www.formula1.com/en/results/2024/races). To contextualize the race times, we draw on molar size data for a large series of marsupials (superfamily Macropodoidea) from Couzens et al. (2016) (see Tables S5 and S6 in the supplementary information of their paper for molar occlusal areas and sample composition description).

To assess these, as in most studies that examine the ICM (e.g., Polly 2007), we plot against each other the sizes of *M*_2_ and *M*_3_, both standardized by the size of *M*_1_. The expectation of the ICM is given by a line with a slope of 2 and a y-intercept of -1. Such graphs conventionally produce a scatter of relative sizes for *M*_2_ and *M*_3_ that are intuited to conform to the expected slope (e.g., Figure 1 in Carter and Worthington 2016, *cf*. Boughner et al. 2021). We note that, using this method, the ICM expectation is met “intuitively” because formal tests of fit using appropriate statistical criteria are absent from the literature.

As an illustration of the degree to which artifacts of standardization can result in intuitively strong fits of even non-ICM consistent patterns of covariance among traits in comparative context, we use an evolutionary simulation. This simulation consists of generating random directional selection gradients, calculating the evolution of the means for the three traits in response to the selection gradients, and using this change to update the position of the simulated lineage in the morphospace. All evolutionary calculations were performed on the mean-standardized scale and converted to the raw scale when recording a simulated population’s position in the morphospace.

We use four instances of the simulation to illustrate how standardization by a common element can lead to artifactual results that give the impression of a fit to ICM expectations. They are:

1. All traits have equal variance (*σ*^2^ = 0.0015) and are not correlated.

2. All traits have equal variance (*σ*^2^ = 0.0015), robust correlations between adjacent elements (*ρ* = 0.75), and zero correlations between non-adjacent (*ρ* = 0)

3. As in simulation 2 above, except the third element has twice the variance (*σ*^2^ = 0.003).

4. All elements have equal variance (*σ*^2^ = 0.0015), adjacent elements are highly correlated (*ρ* = 0.8, and non-adjacent elements are more moderately correlated (*ρ* = 0.5).

The simulation parameters bound plausible relationships that may exist between the measurements of serialized elements, regardless of the developmental or evolutionary model.

## 3 Results

Through our three sets of analyses, we find that the formal expression of the ICM requires a discontinuity in the morphogenic sequence that the model proposes. That is, the proposed inhibiting influence skips the second position in the series and then re-enters the system on the third element. This has implications for extensions of the ICM to systems of serially occurring traits with many elements and no clear point of origin.

Further, we find that the variational expectations of the ICM depart considerably from the assumptions of the regression and phylogenetic comparative methods used to study the developmental, variational, and evolutionary properties of serially occurring traits. Notably, standardization of elements in a series by a single variable leads to spurious correlations between the elements. Even in cases wherein all elements in a series are uncorrelated, common standardization can induce robust and statistically significant correlations among standardized elements. We show, furthermore, that the application of first-element standardization in cases in which adjacent elements are more highly correlated with one another than they are to non-adjacent elements leads to illusionary patterns of association that strongly resemble predictions of the ICM.

### 3.1 Mathematical, developmental, & variational properties of the ICM

As outlined in the mathematical model (Section 2.1), we applied a path model to Equation 2. Inspection of the path diagram in Figure 2 that expresses the ICM on the raw measurement scale, combined with our examination of the developmental evidence used to support the ICM, demonstrates that the ICM lacks the inhibitory component outlined in its written description (Kavanagh et al. 2007). To the extent that inhibition takes place, it is through the difference in the magnitude of the effects of the two developmental pathways (*ϵ*_1_ and *ϵ*_2_).

Note that *ϵ*_1_, which is described as the inhibitory factor *i* in Kavanagh et al. (2007), does not inhibit the growth of the second element (Fig. 2). On the first-element standardized scale used frequently in the ICM, the only reason there is an appearance of an effect of *ϵ*_1_ on *M*_2_ is because of the choice of standardization. That is, the relationship is a mathematical artifact. To the extent that inhibition has a role in the original equation (Eq. 1), it is because the sum of the index of the teeth *k* increases as teeth are added during development. The order in which teeth are added during development is not allowed to vary in the ICM, even if extensions of the ICM suggest ways in which the number of elements might be increased or truncated along the series.

Figure 2 demonstrates another notable feature of the ICM in that it proposes that a developmental process that is singularly and entirely responsible for the ultimate disposition of the first element has no influence whatsoever on the second element and then re-enters the series as one of two predictors of the third and subsequent elements. Biologically, this would involve the masking, silencing, or suppression of the effect of *ϵ*_1_ then reactivation of its effects in a new inhibitory manner from the third element onward. As this is inherent to the original predictive equation for the ICM, such behavior would require substantial revision to the model that would remove it from the ratcheting mechanism proposed in the ICM.

Additional important insights into the variational properties of the ICM are evident from inspection of the path diagram in Fig. 2. The ICM entails a model with two exogenous variables (*ϵ*_1_ & *ϵ*_2_) with two endogenous variables that are in exact proportion to their effects (*M*_1_ & *M*_2_, respectively) linked to a single wholly endogenous variable (*M*_3_). In either case, the ICM predicts that all variation except random error will be a function of the two exogenous variables. In the complete absence of error, the resulting covariance matrix of the endogenous variables will be singular. In the case of a three-element system, one direction in the morphological space will have no variation (see Roseman and Delezene 2019).

The variational predictions on the first element standardized scale are worth mentioning here. Since the IC system has one less degree of freedom than it does endogenous morphological variables, we expect *ρ*(*M*_2_*/M*_1_, *M*_3_*/M*_1_) = 1 in the absence of error, where *ρ* is the Pearson correlation. Then, by the Cauchy-Schwartz inequality, we expect in cases in which all the conditions specified in the ICM hold and in which we may ignore any sources of error:

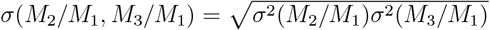

We also expect 2*σ*^2^(*M*_2_*/M*_1_) = *σ*(*M*_2_*/M*_1_, *M*_3_*/M*_1_) in cases without error because the ICM expected slope of the regression of *M*_3_*/M*_1_ onto *M*_2_*/M*_1_ is 2, as noted by Kavanagh et al. (2007).

Since *M*_2_*/M*_1_ and *M*_3_*/M*_1_ share a common denominator, these ratios can be correlated with each other even if all the individual elements that go into their calculation are not correlated (Pearson 1897; Atchley et al. 1976). In a case in which all element sizes have equal variance and are uncorrelated, for instance, we expect *ρ*(*M*_2_*/M*_1_, *M*_3_*/M*_1_) = 0.5. The expected covariance of the errors of *M*_3_*/M*_1_ and *M*_2_*/M*_1_ is simply the mean standardized variance of *M*_1_ (i.e., 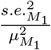). Thus, since *M*_2_/*M*_1_ and *M*_3_/*M*_1_ share a common denominator in *M*_1_, the null expectation for the correlation between the second and third element standardized by the first is not zero. Likewise, when the size of elements occurring in series tend to be more highly correlated with that of their neighboring elements than the size of more distant elements, the expectation for the correlation on the first element standardized scale may be high and produce a regression of *M*_3_*/M*_1_ onto *M*_2_*/M*_1_ that superficially bears a strong resemblance to the ICM expectation, even in the absence of the underlying developmental processes anticipated by the model.

### 3.2 Empirical & simulation examples

Our empirical- and simulation-based examples highlight the effects of the artifactual nature of the apparent goodness of fit of the ICM to empirical data. The empirical example (Figure 3) demonstrates that data from a system that is clearly not governed by a simple ratcheting mechanism, Formula 1 race results, also provide as good a qualitative fit to the ICM as serially occurring morphological features (in this case, the size of the first three molars of marsupials from superfamily Macropodoidea).

**Figure 3.**
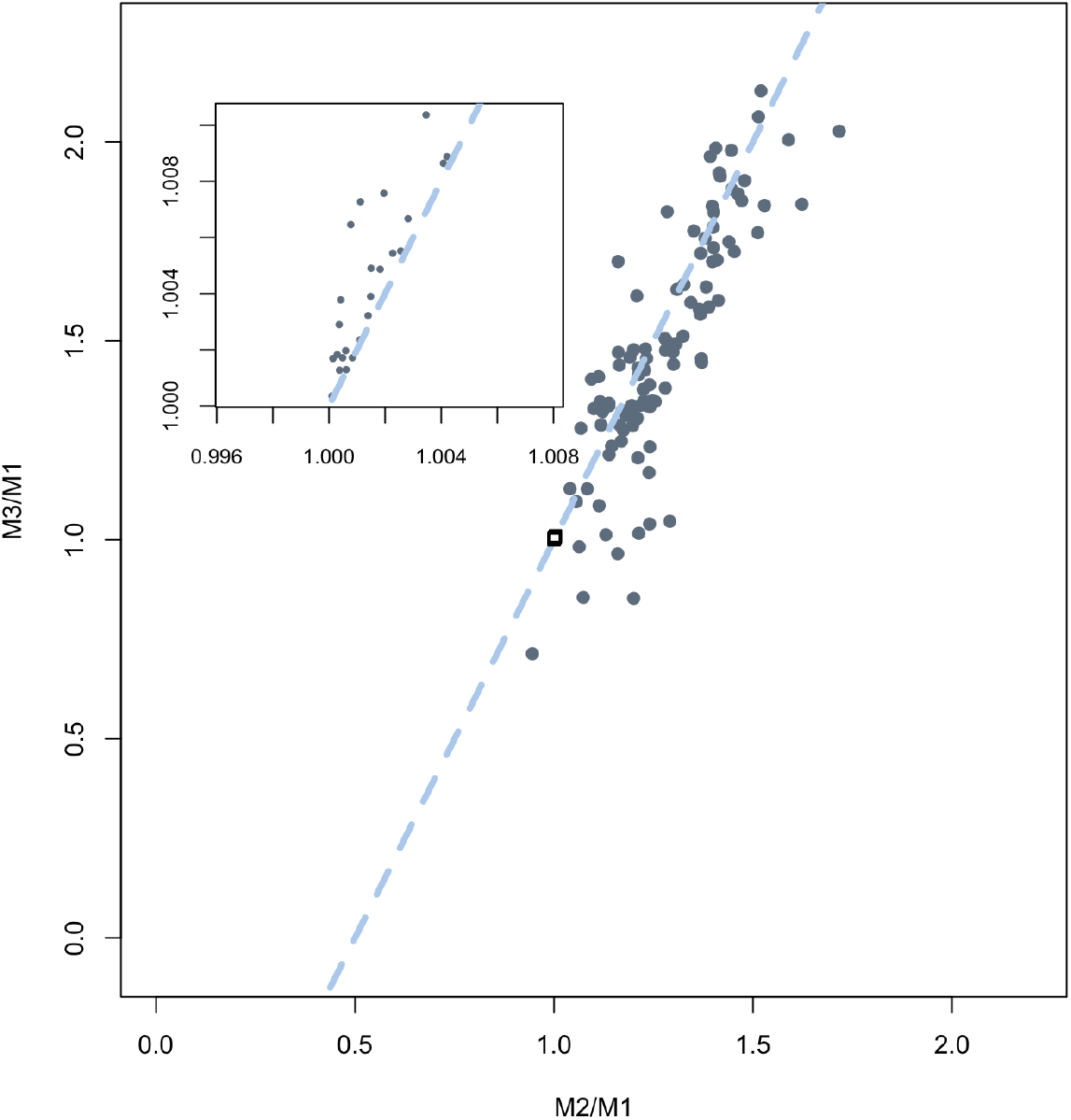
Scatter plots of the the areas of the first three molars of marsupials from superfamily Macropodoidea (Couzens 2016) and of 2024 Formula 1 race finishing times (inset). M2/M1 is the ratio of the second element/finisher and the first element/finisher, M3/M1 the ratio of the third and first elements/finishers, and the dashed lines (intercept -1 and slope 2) are the ICM expected relationship. The inset plot of Formula 1 race times would fit in the square box within the main plot.

Evolutionary simulations of element proportions under different covariance patterns not consistent with ICM nevertheless all yield results that bear a strong superficial resemblance to those from real data presented in the literature. Each example demonstrates that the apparent goodness-of-fit of the ICM to data is driven by artifacts arising from the choice of standardization and not because of any underlying biological phenomenon.

#### 3.2.1 Marsupial teeth & F1 race times

The fit of the marsupial molars to the ICM compares favorably with the subjective fits of similar data reported in other publications (e.g., Polly 2007; Bermúdez de Castro et al. 2021). The inset plot in Fig. 3 depicts the race completion times of the second and third place entries in races held during the 2024 Formula 1 series as standardized by the first place winner in each race. The entirety of the inset fits into the small square close to the *x* = 1, *y* = 1 coordinate in the main plot. Thus, the departure of the points toward the upper left quadrant is not as large as one might intuit given that other studies of the ICM have reported larger departures from the line of expectation as still fitting well within the expectations of the model (see, for example, Machado et al. 2023; *cf*. Boughner et al. 2021). The tendency for Formula 1 relative race times to deviate positively from the ICM prediction line is because race times, by definition, occur in a ranked order from lowest to highest (i.e., *M*_1_ *< M*_2_ *< M*_3_), and so will always plot in the upper left quadrant above the ICM expectation line (see Boughner et al. 2021). Race times tend to be highly correlated because finishing times are closely related to race course distances and the different average speeds obtainable on different courses. Since races tend to be highly competitive, the relative race times are all very close in absolute terms, which restricts them to a very small proportion of the overall plot. That non-biological data on an inorganic phenomenon meet the expectations of the ICM about as well as data from organic systems for which the ICM was designed indicates that an artifact of standardization is likely responsible for both the race time and molar morphological results.

#### 3.2.2 Simulations

In Figure 4, we present the results of simulations of evolution of populations with different among-element covariance structures under random directional selection. None of the covariance matrices we chose in the simulations are consistent with ICM expectations. In all cases in which there are non-zero correlations among elements, outcomes cluster around the line given by the expectation of the ICM. Even when the correlations among the elements are zero, there is still a modest correlation in the simulated evolutionary outcomes on the first element standardized scale. Moreover, even if non-adjacent elements are uncorrelated (*M*_1_ and *M*_3_ are independent), the influence of a common denominator forces data to fit close to the expected slope of 2.

**Figure 4.**
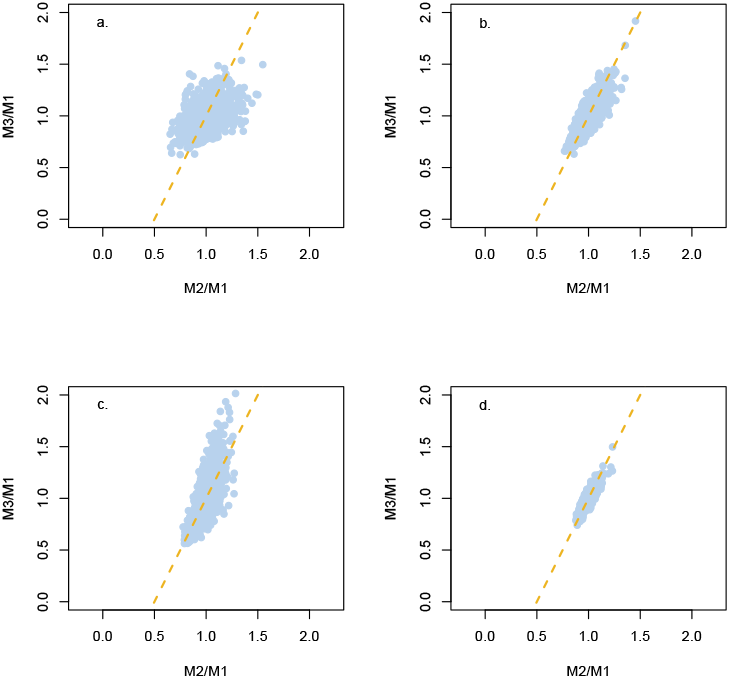
Evolutionary simulations of endogenous variables with different variance and covariance relationships applied to the ICM: a) Uncorrelated endogenous morphological variables with equal variance (*σ*^2^ = 0.0015); b) Positive correlations (*ρ* = 0.75) between adjacent serial elements and zero correlations between non-adjacent elements with equal element variances; c) Conditions as in b except that the third element has twice as much variance as either the first or second elements; d) High correlations between adjacent elements (*ρ* = 0.8) and moderate correlations between non-adjacent elements (*ρ* = 0.5) with uniform variances.

### 4 Discussion

The inhibitory cascade model is an effort to make connections across levels of organization and time scales to unify developmental biology, evolutionary genetics, and macroevolution into a nomothetic theoretical framework. We have demonstrated here that the ICM’s verbal and mathematical descriptions do not correspond well to one another and the choice of standardization artifactually creates patterns that mimic the biological predictions of the model. That the ICM fails empirically is a testament to the fact that it made predictions that are precise enough to reject with some certainty so long as we evaluate them on the appropriate scale of measurement, levels of organization, time scale, and using the right theory (Houle et al. 2011). While the morphologist’s love for the form of life and the developmental biologist’s affection for the dynamics of the assembly of form are vital components of any biology, they are often sidelined in favor of explanations that reduce organisms to parts, sequences, and concentrations of molecules (Auerbach et al. 2023). However, both the holistic appreciation for the organism and the impulse to explain life in terms of ever-smaller phenomena provide an insufficient basis for the building a development-variation-evolution theoretical chain and applying it to biological problems.

Even had we found clear support for genetic sources of inhibition of tooth growth from adjacent teeth in the developmental literature, for example, extending this to other developmental systems of segmented morphology ignores that the established model applies *to the formation of teeth*, which *develop* in series. Vertebrae and limb elements might occur in series, but they do not develop in a series like teeth (Bénazet and Zeller 2009, Maroto et al. 2012; Towers et al. 2012; Skórzewska et al. 2013; Scaal 2016; Serra et al. 2024). The fact that those systems fit the ICM expectations as well as molars is a strong *prima fascie* indicator that an artifact of standardization is responsible for the apparent good fit. This is made clear by the ICM having as the apparent good a fit to results from a non-organic system (Formula 1 race results) as it is to molar development, variation, and evolution. This should be cause for genuine alarm that something is amiss.

A key lesson we can draw from the considerable literature evaluating the ICM in different contexts is the impossibility of discerning the identity and action of biological processes from patterns of variation within populations or from among-group variation on evolutionary time scales on their own. For any pattern of covariance within groups at a given time, there are many ways in which the effects of innumerable small genetic and environmental influences might act on developmental processes to produce that pattern (Hallgrímsson et al. 2012; Hallgrímsson et al. 2023). Similarly, different evolutionary processes and histories may lead to similar patterns of among-group differences. Only a combination of explicit models of development and evolution on one hand and independent predictors of variation and evolution derived from different levels of organization on the other (i.e., genotypic, gene expression, growth rates, strength and patterning of selection, phylogenetic relatedness, effective population sizes, etc.) permits us to make justifiable inferences about the causes of variation (Cheverud 2007; Gompel and Prud’homme 2009; Hallgrímsson et al. 2009; Bookstein 2016).

### 4.1 Our results explain why the ICM failed in other studies

The first indicator of a problem was when data on the sizes of elements from limbs, vertebrae, and other features that *occur*, but do not *develop*, in series fit the ICM expectation as well as mammalian molars for which the developmental component of the model was first proposed. As we state above, the developmental processes underlying molar, limb segment, and vertebral variation take on vastly different forms (e.g., Bénazet and Zeller 2009, Towers et al. 2012; Serra et al. 2024). There is no empirical warrant for us to suppose that a simple two-variable model will describe any one of them, much less all of them.

Many studies made good use of the fact that the ICM can be used to produce clear and testable predictions to identify where the ICM failed to explain morphological change and variation. Using well-known properties of molar development and noting the propensity for many humans not to develop a third molar, Bermúdez de Castro et al. (2021) took advantage of the ICM prediction that the third element would fail to develop in cases in which the first element was much larger than the second due to strong inhibition. Using a large sample of human molar sizes, they demonstrated that this was not the case. This makes sense in light of our demonstration that the actual causal structure of the ICM does not contain inhibition as originally proposed by Kavanagh et al. Similarly, Boughner et al. (2021) found thirteen patterns of molar sizes in humans, and pointed to the variety of patterns reported in other primates as evidence for an array of variational patterns in molar size beyond the ICM.

Carter and Worthington (2016) noted that, if the ICM holds, all of the elements distal to the origin of growth in the series are influenced only by effects that are apparent in full on earlier elements. As a result, the partial correlation between a proximal element and an element further down the series will be zero when conditioned on an intermediate element. In doing so, they expressed the ICM in terms of the language of directional acyclic graphs (DAGs), which is a staple of causal analysis both in the classical (Wright 1921) and modern (Pearl 2009) senses. In doing so, they provided a rough first description of how biological information ought to flow through an ICM-compatible system. The path analysis in our Fig. 2 drew inspiration from their treatment of the problem.

In combination with other analyses of the ICM at different levels of organization, we think that there is effectively no evidence in favor of an IC having much of any role in structuring the within- and among-group variation observed in molar teeth. The evidence that an IC governs the developmental process itself is only marginally better.

Taking a step back from the assumption that molar development is governed by an IC mechanism, Hlusko et al. (2016) resisted the temptation to produce a *post hoc* modification to the ICM to make the pattern fit. Rather, they exercised epistemic humility and declined to make any reaching developmental claims about how variation was generated. Instead, they focused on the relative performance of the trait combinations when it came to recovering phylogenetic signal among groups. Given the data with which they working and time-scale on which their problem was located, this developmentally agnostic approach seems to have been the responsible position.

### 4.2 The specter of artifact in spurious correlations

As we demonstrated, the apparent goodness of fit based on the visual inspection of regressions of correlated variables that share a common denominator is attributable, in large part, to the artefactual correlations induced by the choice of standardization. This problem prompted the original use of the term “spurious correlation,” which was coined by Karl Pearson in 1897. The issue has been revisited periodically in the last century (e.g., Smith 1969; Atchley et al. 1976; Albrecht 1978; Atchley and Anderson 1978; Berges 1997), but the problem keeps rising from the grave and shambling about the literature. Given the tendency for ratios of correlated characteristics to cause analytical and interpretive mischief, it is well past time to put their use to rest.

### 4.3 Conclusion

The ICM has been a prominent attempt to address the development-variation-evolution theoretical chain. It has been a considerable advance to evolutionary developmental biology in that it made precise predictions about how a serially organized and developing set of features might vary and evolve, rather than being purely descriptive. However, as we show, its apparent goodness of fit is a product of artifacts arising from choices of standardization. Moreover, the ICM has important gaps in its mathematical and developmental formulations. Notably, the mathematical rendition of the ICM does not map well onto the description of the development.

## Supporting information

R script for simulation

## 5 Data Availability

All data used in our empirical analyses are available from the sources cited in the Materials and Methods. The R script for evolutionary simulations is available as a supplementary file.

## 6 Author contributions

B.M.A. and C.C.R. conceptualized the original research questions. B.M.A. performed the re-analysis of the developmental framework and worked on the mathematical argument. C.C.R. worked on the mathematical argument, wrote the code for the evolutionary simulation, and performed the empirical analyses. Both B.M.A. and C.C.R. wrote, reviewed, and edited the manuscript.

## 7 Funding

This project was partially supported by a University of Illinois LAS COVID Revitalization Fellowship to C.C.R.

## 8 Conflicts of interest

The authors declare no conflict of interest.

## 9 Acknowledgments

Thanks to Mark Hubbe and Campbell Rolian for helpful discussions about our findings. Leslea Hlusko, Anna Hardin, Mark Grabowski, Arthur Thiebaut, and Mario Modesto Mata provided invaluable feedback on a workshop presentation of a version of this paper. We thank the editor and anonymous reviewers for their comments and suggestions.

## Notes

### Competing Interest Statement

The authors have declared no competing interest.

