## Supplementary material for "The inhibitory cascade model and evolution in segmentally organized tissues": R script for simulation

###################
#
#Date: 20251012 
#
#The inhibitory cascade model and evolution in segmentally organized tissues.
#
#Evolutionary simulation demonstrating artifacts of standardization in ICM comparative analyses 
#
#Arguments:
#
#G  A 3X3 genetic covariance matrix of serially occuring elements on the mean standardized scale
#
#mu A vector of trait means
#
#gen The number of generations of evolution to simulate
#
#Value:
#
#A matrix of first element standardardized second and third elements in a series
#
####################


mouse.sim <- function(G,mu,gen){
  
  #Deploying a matrix for output
  
  out <- matrix(NA,1000,2)
  
  #Iterating over individual simulations.
  
  for(i in 1:1000){
    
    #Updating the means for a new generation of slection.
    
    mu.j <- mu

    #Iterating over generations
    
    for(j in 1:gen){
      
      #Random selection gradient with mean 0 and variance 1. 
      #Gradients are not normalized to length 1
      
      B <- rnorm(3)*mu.j
      
      #Calculate response to selection
      
      delta.z <- G%*%B
      
      #recalculate on raw scale
      
      delta.z <- delta.z * mu.j
      
      #Calculate new position in morphospace
      
      mu.j <- as.vector(mu.j + delta.z)
      
      #The following is included to ensure that all values remain positive.
      
      mu.j[mu.j<=0] <- 0.0001

    }
    
    #Calculate ICM-typical ratios
    
    out[i,1] <- mu.j[2]/mu.j[1]
    
    out[i,2] <- mu.j[3]/mu.j[1]
    
  }
  
  return(out)
  
}
